# Individualized System for Augmenting Ventilator Efficacy (iSAVE): A Rapidly deployable system to expand ventilator capacity

**DOI:** 10.1101/2020.03.28.012617

**Authors:** Shriya Srinivasan, Khalil B Ramadi, Francesco Vicario, Declan Gwynne, Alison Hayward, Robert Langer, Joseph J. Frassica, Rebecca M. Baron, Giovanni Traverso

**Affiliations:** Department of Mechanical Engineering, Massachusetts Institute of Technology, Cambridge, MA 02139, USA; Division of Gastroenterology, Hepatology and Endoscopy, Brigham and Women’s Hospital, Harvard Medical School, Boston, MA 02115, USA; David H. Koch Institute for Integrative Cancer Research, Massachusetts Institute of Technology, Cambridge, MA 02139, USA; Division of Comparative Medicine, Massachusetts Institute of Technology, Cambridge, MA 02139, USA; Department of Chemical Engineering, Massachusetts Institute of Technology, Cambridge, MA 02139, USA; Institute for Medical Engineering and Science, Massachusetts Institute of Technology, Cambridge, MA 02139, USA; Philips Research North America, Cambridge, MA 02141, USA; Division of Pulmonary and Critical Care Medicine, Brigham and Women’s Hospital, Harvard Medical School, Boston, MA 02115, USA

## Abstract

The COVID-19 pandemic is overwhelming healthcare systems worldwide. A significant portion of COVID-19 patients develop pneumonia and acute respiratory distress syndrome (ARDS), necessitating ventilator support. Some health systems do not have the capacity to accommodate this surge in ventilator demand, leading to shortages and inevitable mortality. Some clinicians have, of necessity, jerry-rigged ventilators to support multiple patients, but these devices lack protected air streams or individualized controls for each patient. Moreover, some have not been tested under conditions of ARDS. We have developed the Individualized System for Augmenting Ventilator Efficacy (iSAVE), a rapidly deployable platform to more safely use a single ventilator to simultaneously support multiple critically-ill patients. The iSAVE enables patient-specific volume and pressure control and incorporates safety features to mitigate cross-contamination between patients and flow changes due to patient interdependencies within the respiratory circuit. Here we demonstrate through simulated and *in vivo* pig evaluation the capacity of the iSAVE to support a range of respiratory clinical states. By leveraging off-the-shelf components that are readily available to intensive care unit (ICU) caregivers, the iSAVE could potentially be translated for human application to expand the ventilation capacity of hospitals using existing ventilators, minimizing the need to procure additional ventilators.

## Introduction

### The need for ventilation

The COVID-19 pandemic has severely outpaced the capabilities of healthcare systems worldwide to procure, produce, or adapt the available tools (*1*). The greatest cause of mortality in COVID-19 patients is the development of respiratory failure due to acute respiratory distress syndrome (ARDS). ARDS is ideally managed through intubation and lung-protective ventilation using intubation (*2, 3*). In particular, it is reported that ARDS patients benefit from early intubation, and that early intervention protects healthcare workers from exposure to aerosolized virus from high flow nasal oxygen and non-invasive ventilator machines. However, the volume of patients susceptible to ARDS secondary to COVID-19 is much greater than the number of ventilators available at most hospitals and clinics, as has been seen in Italy, Iran, and China as of March 2020 and more recently in New York City, leading to extremely difficult triage decisions (*4*). Ventilators are vital life-supporting equipment for various conditions treated in ICUs, including chronic pulmonary disorders and surgical emergencies, further limiting their availability.

### Global shortage of ventilators

The most recent publicly available data (2010) reports that approximately 62,000 full-featured mechanical ventilators are available, with an additional 12,700 stockpiled in the CDC strategic national stockpile (SNS)^5^. The demand for ventilators during the COVID-19 pandemic is predicted to significantly exceed the current supply, with an estimated shortage of approximately 960,000 ventilators (*5, 6*). It is infeasible for many hospitals to acquire and maintain this surge capacity of ventilators, given their high cost and temporal demand.

### Approaches to expand ventilator accessibility and supply

Low cost ventilators have been developed to address the cost barrier associated with the acquisition of new ventilators at large scale. While these low-cost designs may represent important innovations, their manufacturing and deployment relies on supply, assembly, and distribution chains that are currently, and may be increasingly, disrupted by the COVID-19 pandemic. Furthermore, the use of new ventilator designs has raised safety concerns, as clinical staff would need to operate unfamiliar technology. Given both supply and implementation hurdles that may delay rapid deployment of low-cost ventilators, other strategies warrant consideration.

An alternative strategy previously proposed is the splitting of one ventilator to multiple patients, which has been performed in a few emergency cases (*7*). By utilizing readily available tubing and ventilatory equipment, this approach is immediately scalable and permits use of ventilators already familiar to clinicians. Multiplexing ventilation involves connecting multiple outflow tracts to the ventilator to dividing the flow amongst the patients (*8*). In this configuration, the compliance and resistance parameters of each patient’s pulmonary systems become part of the same circuit. This yields patient interdependence, which poses the following major safety concerns. 1) Independent control of volume and pressure to each patient is not possible, which is important for lung-protective ventilation standard of care for ARDS. 2) Changes in one patient’s condition (clinical improvement or deterioration) results in an automatic change to ventilation of other patients. 3) Sudden events such as pneumothorax, tube occlusion, or disconnection of an endotracheal tube, causes harmful imbalances of ventilation that are potentially deleterious for other patients. 4) Cross contamination of airborne pathogens across channels can occur. 5) Alarm monitoring becomes challenging due to a complex circuit configuration. For these reasons, splitting ventilators has not gained widespread acceptance. In fact, a number of medical associations, including the American Association for Respiratory Care (AARC), issued a joint statement explicitly advising clinicians against the sharing of mechanical ventilators with current equipment, despite the dire need with the current COVID-19 pandemic (*9*). Our work aims to overcome these limitations.

### Conceptualization of a patient-specific ventilation expansion system

To address the need to expand ventilator capacity whilst incorporating the translational constraints associated with a rapidly developing pandemic such as COVID-19, we have developed the Individualized System for Augmenting Ventilator Efficacy (iSAVE). This system repurposes existing flow regulation valves in the medical industry to utilize a single ventilator to provide personalized support to at least two patients. The iSAVE’s ventilation circuit enables independent control of volume and pressure for each patient and incorporates safety measures to accommodate sudden patient deterioration and cross contamination. Here, we describe the design and validation of the iSAVE through benchtop and *in vivo* tests splitting a single ventilator to two patients. We hypothesize that our system will be capable of maintaining desired ventilation parameters to each patient amidst static and dynamic changes in resistance and compliance. We simulate several clinical scenarios, focusing on those relevant to the management of ARDS, and validate the safety mechanisms of the iSAVE. Leveraging off-the-shelf components, the iSAVE can facilitate a rapid expansion of the ventilation capacity of hospitals.

## Results

### Design of the System for Augmenting Ventilation Efficacy (iSAVE)

Ventilators aim to achieve two main goals, oxygenation and ventilation. Modern ventilators enable the control of various parameters including tidal volume (V_T_), inspiratory pressure, positive end expiratory pressure (PEEP), fractional oxygen concentrations (FiO_2_), and respiratory rate. Patients with ARDS are usually ventilated using low V_T_ ventilation and high PEEP to limit lung injury (lung-protective ventilation), which has been shown to improve patient mortality.^11^ For lung-protective ventilation using volume-control, V_T_, PEEP, and respiratory rate are carefully set based on individual patient parameters while inspiratory pressure varies. Alternatively, pressure control ventilation can be used, in which pressure applied to the airway is constant and V_T_ varies.

The iSAVE utilizes a series of valves and flow regulators in parallel limbs to effectively maintain the desired volume for each patient (Figure 1A-C, Supplemental Table 2). Here, we describe the overall design, components and safety features of the iSAVE.

**Figure 1.**
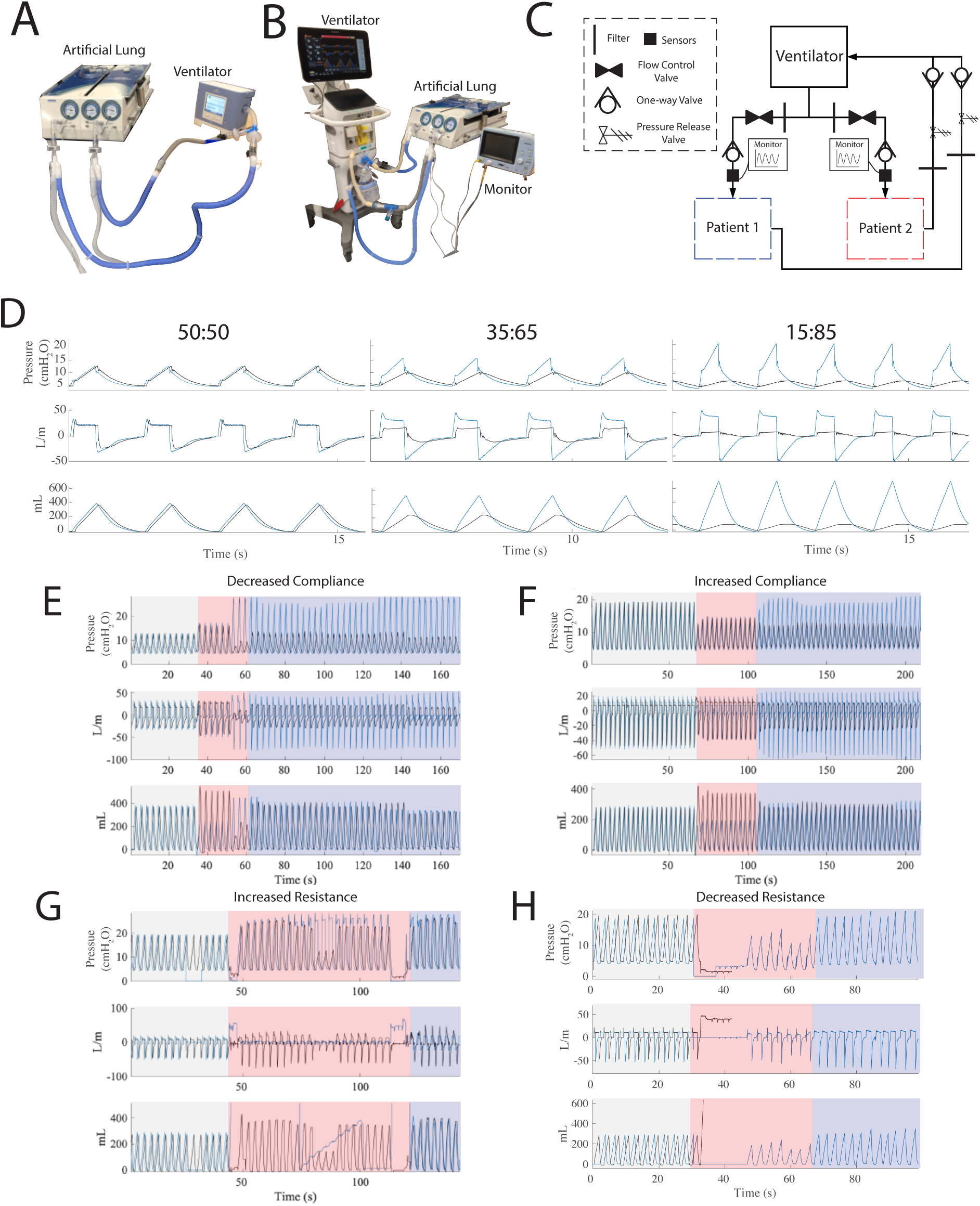
Individualized System for Augmenting Ventilation Efficacy (iSAVE) Design and Validation. A) iSAVE with open-circuit (single limb) ventilator consists of filters and flow valves connected in series on each channel. Pictured are two channels connected to a test lung apparatus for benchtop validation. B) iSAVE with closed-circuit (dual limb) ventilator employing 2 channels connected to a test lung apparatus for benchtop validation. The Philips NM3 respiratory monitor is used to monitor the pressure, flow, and CO_2_ in each patient circuit. C) Circuit diagram to implement the iSAVE on closed-circuit ventilators. D) Variable tidal volumes are delivered through the system. E) Left: Simulation of decreased compliance in one lung (blue) and (Right) increased compliance in one lung (black) G) Simulation of increased resistance in one lung (blue) and (Right) decreased resistance in one lung (black).

In open-circuit ventilators (Figure 1A, Supplemental Figure 1), Y or T connectors are used to provide individual channels for each patient. Each channel incorporates a low resistance bacteria/viral respiratory filter, a flow control valve, and a set of sensors in series prior to the endotracheal tube. The expiratory limb contains another filter and is connected to a Whisper Swivel that serves as the expiratory flow port. See Supplemental Tables 1 and 2, which provide a concise list and description of the medical names for equivalent valves.

For closed-circuit ventilators (Figure 1B-C), Y or T connectors are utilized to provide individual inspiratory channels for each patient. Each inspiratory channel consists of a filter, flow control valve, a one-way flow valve and a set of sensors in series prior to the endotracheal tube. The expiratory limb consists of a filter, pressure release valve (a PEEP valve can be used for this purpose), one-way valve and filter prior to connection to Y or T connectors which are routed back to the ventilator.

The flow control valve is used to allocate the appropriate V_T_ to each patient. Bacterial/viral filters on each limb prevent cross-contamination between individual patient circuits. The filters also filter expired gas before release into the room through the pressure release valve, thus limiting pathogen exposure to healthcare workers. The one-way valves prevent backflow and mitigates over-distention in cases of rapid flow change. PEEP valves enable the release of excess pressure that may be caused by changes in the flow to each limb. Pressure and flow sensors and a mainstream capnometer are positioned in series with the inspiratory line for set up, titration and monitoring.

Data from these sensors can be visualized on separate patient monitors or on the ventilator. During the initial set up, the respiratory rate, PEEP, FiO_2_ and inspiration:expiration time, and the sum of the V_T_ for both patients is set on the ventilator. Then, the flow control valves can be titrated to provide the desired volume to each patient. The exhaled volume and minute ventilation alarms on the ventilator are set based on the sum of the exhaled tidal volumes from both patients. This alarm will enable the ventilator to notify clinicians in the case of any sudden changes in the volume delivered to each patient (e.g., shunt, occlusion or, disconnection of the endotracheal tube). PEEP valves in each patient’s circuit should also be set to the maximum pressure allowable for that patient.

### Benchtop testing of the iSAVE

#### Delivery of Variable Patient Volumes

Given its immediate application to support patients in ARDS who are normally ventilated through volume-controlled ventilation, we validated the ability of the system to deliver variable volumes to each patient.

The iSAVE connected a ventilator delivering a V_T_ of 800 mL to two test lungs with healthy tissue characteristics (compliance (C) = 50 cm H_2_O and resistance (R) = 5 cmH_2_O/L/s) and diseased tissue characteristics with low compliance (C = 20 cmH_2_O, R = 5 cmH_2_O/L/s). By titrating the flow control valve, we were able to achieve variable V_T_ delivery spanning splitting ratios from 50:50, to 15:85 (Figure 1D). However, at ratios above 20:80, pressure exceeded 40 cm H_2_O, which is beyond the therapeutic range.

#### Accommodation to static compliance changes

Ventilator settings and respiratory system properties for ARDS patients can vary significantly and evolve rapidly through the course of the disease and recovery. We thus tested the ability of the iSAVE to compensate for static changes in compliance and resistance of one lung while minimizing effects on the other patient. For all tests, we 1) measured the baseline ventilation values (graphed in gray), 2) performed an intervention (changing compliance or resistance, graphed in red), 3) noted any safety features or alarms that were activated and 4) titrated valves to return to the baseline values (within 5% error) (graphed in purple).

In the first test, we decreased the compliance of one of the lungs (blue) (C: 50 mL/cm H_2_O to 15 mL/cm H_2_O), simulating the parameters characteristic of ARDS (Figure 1E). Flow was quickly diverted, resulting in a disproportionate volume being delivered to the healthy lung (C: 50 mL/cmH_2_O). The PEEP valve released excess volume during the period of titration, preventing overdistention of the healthy lung. The desired volume of 400 mL/lung was restored by titrating the flow control valve. We then started with two lungs simulating ARDS (C = 20 mL/cm H_2_O, R = 5 cm H_2_O/L/s) and increased the compliance of one lung (C: 15 mL/cm H_2_O to 50 mL/cm H_2_O black), which created a shift in the volumes delivered. Adjustment of the flow control valve restored the desired flow to both lungs (Figure 1F) in less than 13 breaths.

#### Accommodation to static resistance changes

We then performed ventilation (300 mL/lung) under high resistances, simulating the physiology characteristic of the comorbidities commonly associated with ARDS, including asthma, COPD/emphysema, and presence of viscous airway secretions (commonly reported in COVID-19 infections). With both lungs simulating ARDS (C = 20 mL/cm H_2_O, R = 5 cmH_2_O/L/s), the resistance of one lung was increased ten-fold (R 5 cm H_2_O/L/s to 50 cm H_2_O/L/s, black), causing a drastic reduction in the flow to the lung. Titration of the valve along with an increase in the inspiratory time enabled the desired flow while maintaining lower pressures (Figure 1G).

#### Safety measures

While subacute changes in resistance and compliance can be accommodated, it is vital that the iSAVE provide alerts in response to acute changes for patient safety. The ventilator alarm was set to detect changes in the overall expiratory volume. We mechanically occluded tubing of one lung to simulate an instantaneous change in resistance, which created a reduction of flow in one channel and spike in pressures/volumes of the other (Supplemental Figure 2). This instantaneously caused the ventilator alarm to activate. We also simulated the loss of the endotracheal tube to one lung, which causes a leak in the system yielding minimized flow to either lung. This activated the main ventilator leak alarm. We closed the valve to the disconnected lung and titrated the other valve to deliver the desired volume (Figure 1H). This process can be additionally utilized to remove a patient from the ventilator (in cases such as cardiac arrest or weaning from the ventilator) without leak into the room, aerosolizing virus and exposing healthcare workers.

The same benchtop tests were performed with an open-circuit ventilator, yielding identical results. See Supplemental Materials for details.

### In vivo testing

We performed ventilation of a large animal (Yorkshire Swine, 70kg) alongside an artificial lung, delivering a total V_T_ = 500mL from the ventilator (Figure 2A). The swine possesses a 5-6L capacity, similar to human lungs and serves as a good model for this testing. Oxygen saturation, ETCO_2_, RR, and V_T_ delivered to the swine and test lungs were monitored. The iSAVE was able to deliver variable volumes to each channel. Ratios of 50:50, 40:60, and 30:70 are presented in Figure 2B.

**Figure 2.**
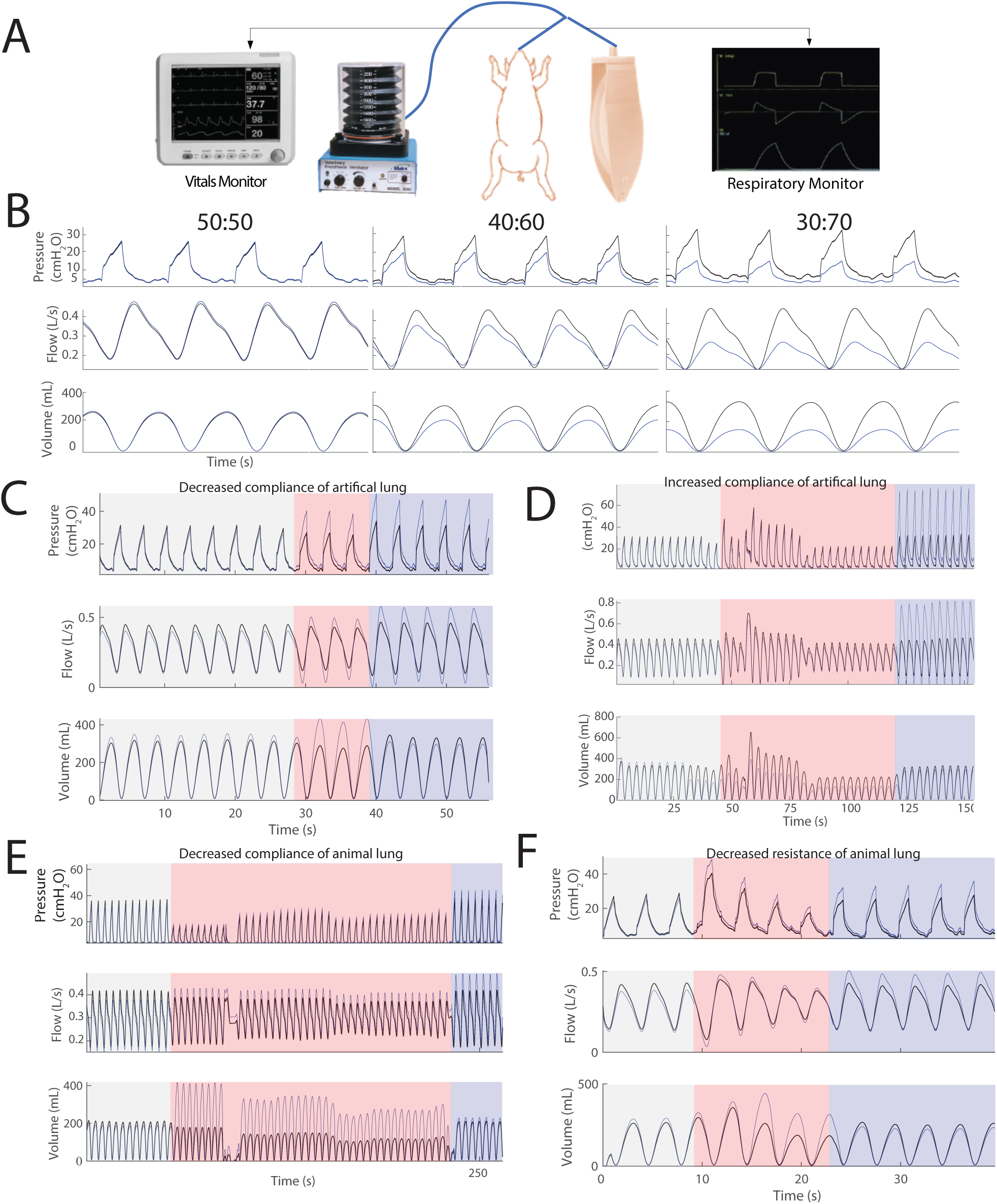
*In Vivo* Validation of the iSAVE) A) Set up for *in vivo* testing consisted of a single animal ventilator supporting a 70kg pig (5-6L lung capacity) and artificial test lung (1L capacity). B) Variable V_T_s are delivered to the pig and artificial lung. C) Simulation of decreased compliance in the artificial lung (black) and D) increased compliance in the artificial lung (black). E) Simulation of decreased compliance of the animal lung following 100mL of saline infusion (black) F) Decreased resistance of the artificial lung (black).

#### Accommodation to static compliance changes

V_T_ was equally distributed from the ventilator to each branch (600 mL total, 300 mL per branch). The compliance of the artificial lung was decreased (black, C 120 mL/cm H_2_O to 60 mL/cm H_2_O), resulting in a greater allocation of volume (∼100mL) to the animal (blue) (Figure 2C).

By titrating flow with the valve, we restored V_T_ distribution within 3 breaths. During the course of recovery in ARDS, lung compliance increases. Thus, we tested a range of such scenarios: C 20 mL/cm H_2_O to C 50, 60 and 120 mL/cm H_2_O. In Figure 2D, we present the most extreme case to test the iSAVE’s capacity to perform adequate accommodation (black, C: mL/cm H_2_O to 120 mL/cm H_2_O). This immediately shunted flow away from the higher resistance in the animal, towards the less resistant artificial lung. Valve adjustment restored the desired flow to the animal’s lungs and artificial lung. The animal’s lungs were filled with 100mL of saline, effectively decreasing their compliance and resulting in a lower V_T_. With repeated breaths, flow was further diminished despite efforts by the automatic adjustments of the ventilator in increasing the pressure (Figure 2E). Titration of the flow valve enabled a restoration of the desired volume. Throughout these tests, ET PCO_2_ and SpO_2_ were maintained between 38-41mmHg and 91-98%, respectively, at baseline and after titration.

To simulate scenarios such as tube clogging and aspiration, the endotracheal tube of the artificial lung was mechanically obstructed, immediately increasing flow to the animal by 30% (Figure 2F). This was quickly resolved by closing the valve to that circuit and adjusting flow to the animal.

## Discussion

In this study, we demonstrate that the potential for iSAVE to deliver multiplexed and personalized ventilation through a system of valves and sensors, with embedded safety measures. Our benchtop and *in vivo* testing simulated the characteristics of patients with ARDS as well as clinical scenarios associated with multiplexing ventilation. The results show that the system enables patient-specific titration of V_T,_ inspiratory pressure, and PEEP, significantly amplifying the capacity of a single ventilator. In a set of scenarios mimicking the clinical evolution of patients with ARDS and sudden events that could jeopardize the ventilation of other patients, the system is able to handle circuit dependencies and deliver the desired therapeutic parameters to each channel. Our approach employs valves and sensors to similarly prevent life-threatening shunting of flow away from diseased patients to relatively healthier patients. The positioning of circuit components further overcomes issues of backflow, contamination, and higher pressures. We also validated that obstruction or disconnection of a tube will activate pressure or volume-based alarms in the ventilator. These results address the critical challenges expressed by the medical community implicated in splitting ventilation (*9*).

In addition to the components described and demonstrated here, CO_2_ sensors could be easily incorporated into the setup for personalized monitoring of inspired and expired air in each patient circuit. A whistle ring could be added to the pressure release valve to provide an additional auditory alarm when a change occurs in the system causing high pressure in one segment (See Supplemental Figure 3). Future iterations could incorporate closed-loop control, directly modulating flow based on flow meter readings and valves to allow for independent control of RR and FiO_2_ in each patient circuit. Although, this system, as described, permits individualized minute ventilation through alteration in tidal volume for each patient. Finally, the capacity to derive plateau pressures can be built into the circuit. In theory, this system can accommodate more than two patients, although this may present significant practical challenges. Please see Supplemental Note 1 for more details.

The open-circuit system is less complex than the closed-circuit system, requiring fewer parts and having fewer backflow issues. While the expired air is filtered before being vented through the expiratory limb, the high aerosolization risk of a virus such as COVID-19 may present a contamination risk to other patients and healthcare personnel. Thus, a closed-circuit system may be preferred in some settings. However, in settings where multiple patients are cohorted in a single, large space, where healthcare providers continuously wear full personal protective equipment (PPE), the preference for a closed-circuit (dual limb) ventilator may not apply. In these settings it may be preferable to use single limb devices for their simplicity. The iSAVE is intended to be used in a negative pressure room, or other isolation setting that would not be entered by healthcare workers without PPE.

### Translational Considerations and Clinical Management

The system’s novel components (pressure release valve, flow control valve, one-way valve, pressure sensor, CO_2_ sensor) are commercially available parts that are commonly found in hospitals, lowering the barrier to implementation (See Supplemental Table 2). Alternatively, in a worst-case scenario, these parts can be easily procured from hardware stores (from departments such as plumbing and ventilation) and autoclaved for sterile usage. If standard adaptors do not interface these parts, adaptors can be made from standard piping or through 3D printing.

In the envisioned set up, individual patient monitors (such as the NM3 (Philips)) or similar monitoring equipment integrated with ventilators should be utilized to set the initial conditions of the valves, perform periodic checks, and determine changes in the circuit. If standard sensors are employed, they can also be connected to the ventilator.

Based on the variable V_T_ ratios capable of the iSAVE, matching criteria for patients would not need to be as stringent as other protocols designed to multiplex ventilation without individualized control. Despite the robust nature of the iSAVE, certain ventilatory parameters (RR, FiO_2_, I:E, PEEP) will remain shared amongst patients. To this end, patients must remain sedated to a degree that prevents spontaneous breathing, which would lead to asynchrony in the system. Thus, patients should be matched as closely as possible in terms of degree of illness and ventilatory needs to optimize functioning. We recommend the following types of stratification to match patients when feasible:

1. <u>FiO_2_</u>: Patients requiring FiO_2_ > 60% should be grouped separately from patients requiring FiO_2_ < 60% given common FiO2 settings and a desired threshold to lower FiO2 < 60% to avoid theoretical O_2_ toxicity.
2. <u>Compliance:</u> Patients with a C < 30 cmH_2_0 should be grouped separately from patients with C > 30cmH_2_O.
3. <u>PEEP</u> should be within the range of 5-18 cmH_2_0 and the difference between patients should be minimized to less than 5 cmH_2_0.
4. <u>Driving pressure</u> should be within the range of 5-15 cmH_2_0 and the difference between patients should be minimized to less than 6 cmH_2_0.
5. <u>Difference in height</u> should be minimized to 3-6 inches to ensure relatively similar tidal volume needs.
6. <u>Resistance</u>: Patients presenting with restrictive comorbid diseases (e.g., asthma, emphysema/COPD, bronchiectasis) should be grouped similarly to make circuit dynamics more uniform.

Other considerations include hemodynamic stability, anticipated invasive ventilation time, co-infection and the logistics of space allocation for patients. Ideally, these patients would be ventilated in negative pressure rooms, with their heads being as close as possible to the ventilator. Recovering patients would be transitioned to a standard ventilator when spontaneous breathing becomes viable (*10*).

### Limitations and Path to Translation

Our aim is to ensure that the iSAVE can be reliably implemented and utilized across intensive care clinical settings in cases of ventilator shortages. To this end, further steps must be taken to address current limitations. 1) Due to differences in performance characteristics (mechanism of pressure/flow monitoring) of ICU ventilators, this approach must be tested across a range of ventilators in both volume and pressure control modes. 2) The iSAVE must be evaluated in conditions reflecting the real-life variability of intensive care practice. 3) The procurement and the sterilization process for non-standard components must be addressed. 4) First-in-human trials must be performed to replicate the results from benchtop and animal investigation. Towards facilitating this evaluation by our team and others, we have assembled a list of components (Supplemental Table 1).

### Ethical Considerations

While this system mitigates several challenges associated with splitting ventilation, it has several drawbacks, unknown limitations and should be deployed with careful consideration. In a setting of severe life-threatening shortage of ventilators, ventilating two patients with the potential to save two lives may mitigate the need to triage which patients receive ventilatory support and which do not. This may present greater overall societal value in comparison to potentially only saving one life and allowing the other to pass. Further ethical and policy-based discussions are necessary prior to implementation.

## Conclusion

The iSAVE is a rapidly deployable solution in response to the urgent shortage of mechanical ventilators for respiratory support during the COVID-19 pandemic. In contrast to previously described ventilation splitters, the iSAVE could at least double the number of treated patients while retaining personalized ventilation settings for each individual. Using existing ventilator devices avoids manufacturing delays, and lowers the prohibitive costs of procuring expensive ventilators at large scale. Healthcare systems worldwide could benefit from this as they strive to care for the exponentially increasing volume of COVID-19 patients.

Disclaimer: The authors caution that the iSAVE approach is not a standard of care and should be considered only in the most dire settings where shortages of patient ventilators threaten lives. With further testing and validation, implementation may be classified under the Emergency Use Authorization issued by Food and Drug Administration for ventilators for use in healthcare settings to support patients during the Coronavirus Disease 2019 pandemic (*11*). The techniques and approaches described in this article do not represent a recommendation or alteration in the recommended use of any device that was used in this demonstration and study. The use of any technique described in this article is subject to the clinical judgement of the physicians caring for individual patients.

## Materials & Methods

### Open-circuit ventilator assembly

A Philips Trilogy portable ventilator was utilized as a representative open-circuit ventilator for testing. Standard flex corrugated tubing was utilized to connect two respiratory circuits consisting of a 1) ball valve (EKWB G ¼’’ Nickel, Microcenter), 2) bacterial filter (Main Flow Bacterial/Viral Filter, Teleflex Medical), 3) pressure sensor (TD160D, Biopac Systems), 4) airflow sensor (BSL Medium Airflow Transducer SS11LA and AFT 20, Biopac Systems) and 5) Whisper Swivel valve (passive exhalation port, Philips). This was connected to the expiratory side of a Y-piece to allow for the inclusion of a bacterial/viral filter before venting exhaled gas to the ambient. (see circuit diagram provided in Figure 1A, exhalation filter not illustrated). The airflow and pressure sensors were connected to DA100C differential amplifiers with 1000x gain and processed through the MP160WSW signal processing unit (Biopac Systems) sampling at 2kHz. Volume control mode was employed, and settings were adjusted to deliver the desired V_T_, RR, and PEEP. Artificial linear test lungs (IngMar Medical) were utilized for simulations with the open-circuit ventilator.

### Closed-circuit ventilator assembly

A Philips critical care ventilator was utilized for testing as representative of closed-circuit ventilators typically used in the ICU. The iSAVE is designed to work with virtually any closed-circuit ventilators currently found in ICUs. Y-connectors were connected to the inspiratory and expiratory limbs of the ventilator to enable individual channels for each lung. In the inspiratory limbs, the following components were connected in series: 1) a bacterial/viral filter (Main Flow Bacterial/Viral Filter, Teleflex Medical), 2) ball valve (EKWB G ¼’’ Nickel, Microcenter), 3) one-way valve (a Threshold PEP positive expiratory pressure device, Philips, was used as a surrogate for one-way valves), 4) pressure sensor, 5) flow sensor, and 6) capnostat adaptor. In the expiratory limb, the following components were connected in series: 1) a bacterial/viral filter, 2) a pressure release valve (PEEP valve), and 3) an optional one-way valve prior to connection with the ventilator (Threshold PEP). PTFE tubing was used to adapt the ball valves to the standard 22mm outer diameter corrugated flex tubing. A dual adult lung simulator (Model 5600i, Michigan Instruments) was used to perform simulations with the closed-circuit ventilator. Pressure and flow were displayed and recorded by the Philips NM3 monitors and associated sensors, one for each channel.

### Benchtop testing protocol

1. <u>Check for leaks and alarms:</u> We first tested the ability of the system to perform ventilation without leaks. We then disconnected a lung to ensure the activation of standard alarms on the ventilator.
2. <u>Variable tidal volume production:</u> Starting with both flow valves fully open, we measured the flow and pressure delivered to each lung. Then, we gradually closed one of the valves, measuring the distribution of flow, to map the range of volume distribution capable of the system.
3. <u>Maintenance of V_T_ following static changes to compliance and resistance</u>: Patient interdependencies in the circuit pose a major challenge for the multiplexing approach of ventilation. As a patient degenerates or recovers, changes in tissue compliance and resistance occur. These cause shifts in the overall circuit’s flow dynamics. We modulated the compliance and/or resistance of one test lung, measured the effect of the intervention, and titrated the valve until the baseline parameters were reached. Resistors (Rp5, Rp20, Rp50 and Rp500 Michigan Instruments) were utilized for all benchtop testing. We performed these tests with baseline parameters set to those of 1) a healthy lung (C = 50 cmH_2_0, Rp5) 2) a lung with ARDS (C = 20 cmH_2_0, Rp5-50). While the flow valve was the main variable we modulated, in some cases, the ventilator’s tidal volume or inspiration : expiration ratio was also adjusted. These cases are specifically mentioned in the results section.
4. <u>Maintenance of V_T_ following dynamic changes to compliance and resistance</u>. Sudden deterioration of a patient (blood clot, pneumothorax) or equipment malfunction (kinked endotracheal tube) pose great risk to the balance of ventilation between patients. We tested the ability of the iSAVE to manage these scenarios. We simulated acute changes by disconnecting or clamping tubes and monitoring the ventilator’s response. The flow valves were also titrated to return the circuit to its baseline.

### In vivo testing

Animal experiments were approved by the Committee on Animal Care at the Massachusetts Institute of Technology (MIT). The experiment was performed as part of a terminal procedure on a 70kg female swine (n=1). The iSAVE was connected to a veterinary anesthesia ventilator (Model 200IE, Hallowell EMC) delivering 2% isoflurane in oxygen. One inspiratory circuit was connected to the anesthesia machine delivering gas to the animal while the other inspiratory circuit was connected to an artificial linear test lung (IngMar Medical). Pressure and flow measurements were recorded on the inspiratory limb of the animal. A VetTrends Vital Signs Monitor was utilized to measure SpO_2_, ETCO_2_, RR, and other physiological parameters. 600mL of V_T_ were equally distributed between the animal and test lung. A respiratory rate between 18-20/min was set on the ventilator. We carried out the same tests outlined in the benchtop testing protocol, modulating the parameters of the test lung, to validate the capabilities of the iSAVE to restore the system to baseline. Once the animal was euthanized, to acutely change the compliance of the pig’s lung, we utilized an endoscope (PENTEX) to deliver 50mL of phosphate buffered saline (PBS, Sigma Aldrich) into the left and right bronchi. We then ventilated the animal, aiming to restore tidal volume to the baseline values prior to euthanasia. Approximately 10 minutes later, we delivered an additional 700mL of saline into the right and left bronchi. Ventilation was performed and the valves were titrated to achieve desired flow parameters.

### Data Analysis

Pressure and flow data from the Biopac system was analyzed using MATLAB (2018a). Flow data was integrated every respiratory cycle to derive the flow. In cases where pressure and flow data were not simultaneously obtained from both limbs of the circuit, it was derived based on the settings on the main ventilator and the measurements on one limb.

## Acknowledgements

We want to thank Kevin Wasco, Ken Graap, and the Biopac team for extreme responsiveness for provision of sensor equipment, and Patricia Kritek for helpful respiratory related discussion and guidance. We’d like to thank Aartik Sharma, Joshua George, Jake Wainer, Adam Wentworth, for their assistance in prototyping and/or consultation. This work was funded by in kind services from Philips, discretionary funds from the Department of Mechanical Engineering at MIT and at BWH to G.Traverso.

## Author contributions

S.S.S performed the conceptualization, investigation, data analysis, and writing. A.H. assisted with in vivo investigation. K.R. performed writing. F.V. performed investigation, data curation, and edited the manuscript. D.G assisted with prototyping. R.L, J.F., D.H., R.M.B, and G.T. contributed to conceptualization and editing of the manuscript. All authors approved the manuscript.

## Competing interests

F.V. is an employee of Philips., J.F is an employee of Philips. R.M.B. is part of an Advisory Board for Merck. All other authors report no competing interests related to the work reported here.

## Data and materials availability

All data associated with this study are in the paper or supplementary materials.

