## Supplemental Materials for "Individualized System for Augmenting Ventilator Efficacy (iSAVE): A Rapidly deployable system to expand ventilator capacity"

**Table 1. List of components required for the assembly of the iSAVE**

| **Required Components for Closed-circuit (dual limb) Ventilator** |
|---|
| Per patient:<br>[ ] 2 one-way valves<br>[ ] 1 pressure release valve<br>[ ] 1 flow regulator<br>[ ] bacterial/viral filters<br>[ ] set of pressure/volume sensors<br>[ ] capnostat<br>[ ] respiratory profile monitor |
| Required Components for Open-circuit (single limb) Ventilator |
| Per patient:<br>[ ] 1 flow regulator<br>[ ] 1 bacterial/viral filter<br>[ ] 1 set of pressure/volume sensors (optional)<br>[ ] 1 capnostat (optional)<br>[ ] 1 monitor (optional) |

**Supplemental Table 2. Mechanical components used in the iSAVE and their equivalents, readily available in the medical industry.**

| Mechanical Component | Medical Equivalent |
|---|---|
| One-way valve | Positive expiratory pressure (PEP) threshold device<br><br>Unidirectional resistance/one-way valve<br><br>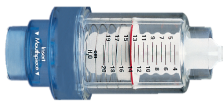 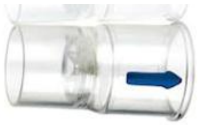 |
| Pressure release valve | Positive end-expiratory pressure (PEEP) valve<br><br>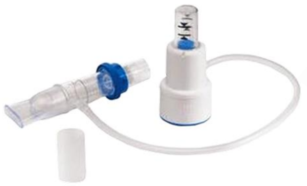 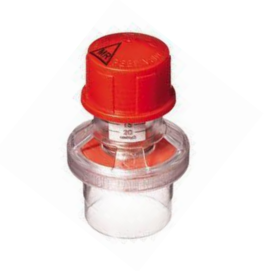 |
| Flow regulator | 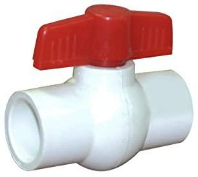 |
| Bacterial/viral filters | 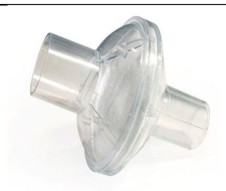 |
| Pressure/volume sensors | 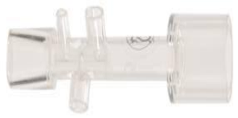 |

| | |
|---|---|
| $CO_2$ sensor | E.g., Capnostat for volumetric capnography (including $ETCO_2$ monitoring) or LoFlo for $ETCO_2$ monitoring 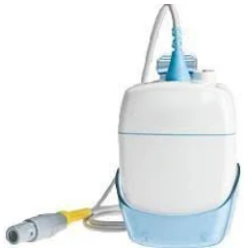 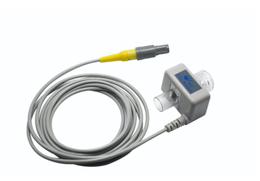 |
| Respiratory profile monitor | 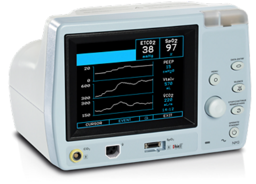 |

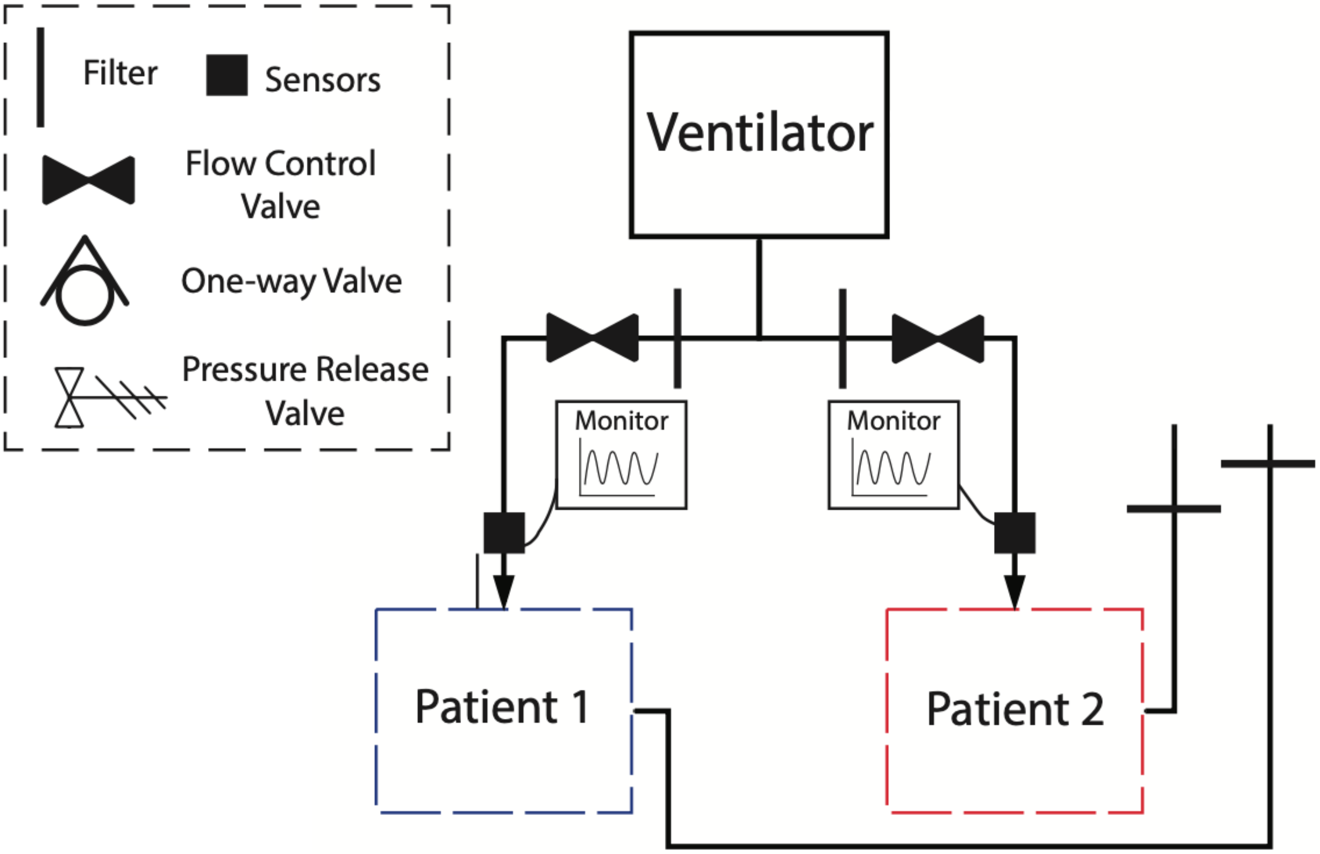


**Supplemental Figure 1.** Circuit diagram to implement the iSAVE  on open-circuit ventilators.

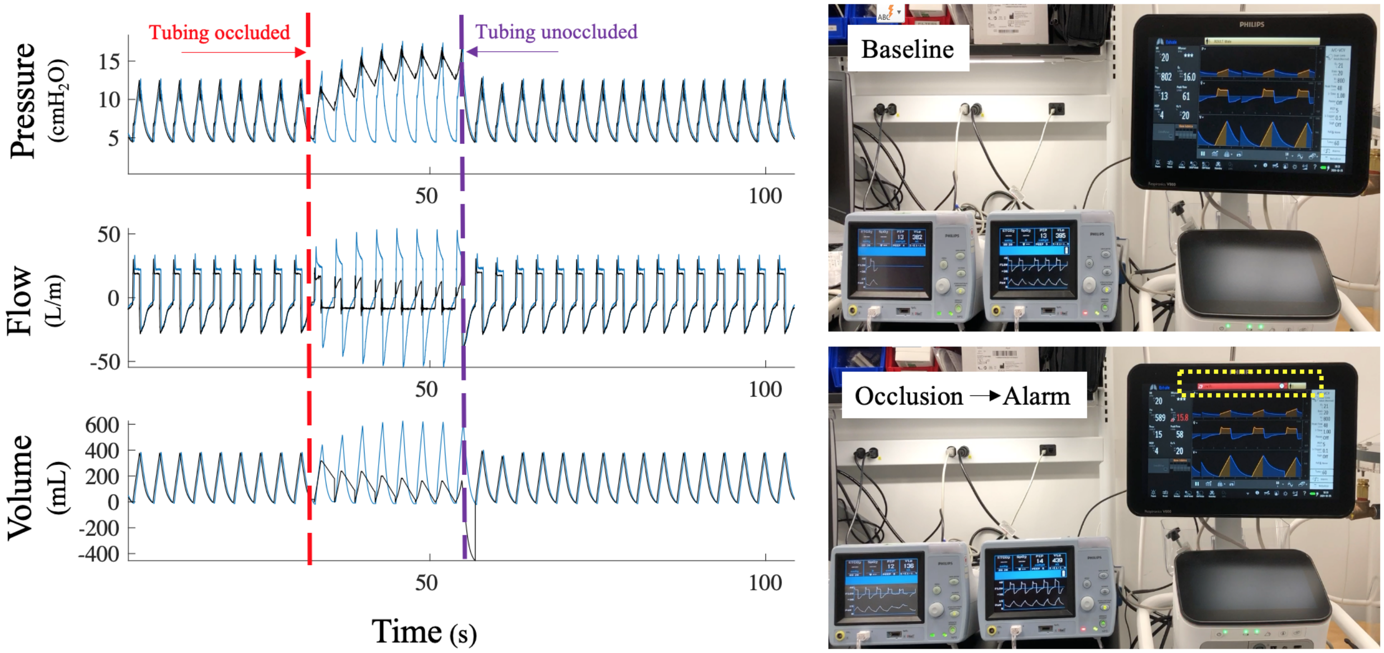


**Supplemental Figure 2.** Tubing of one lung (black) was occluded to simulate the case of sudden events causing a high resistance or complete shunt in one limb. The ventilator's alarm immediately activated. This was resolved by removing the occlusion.

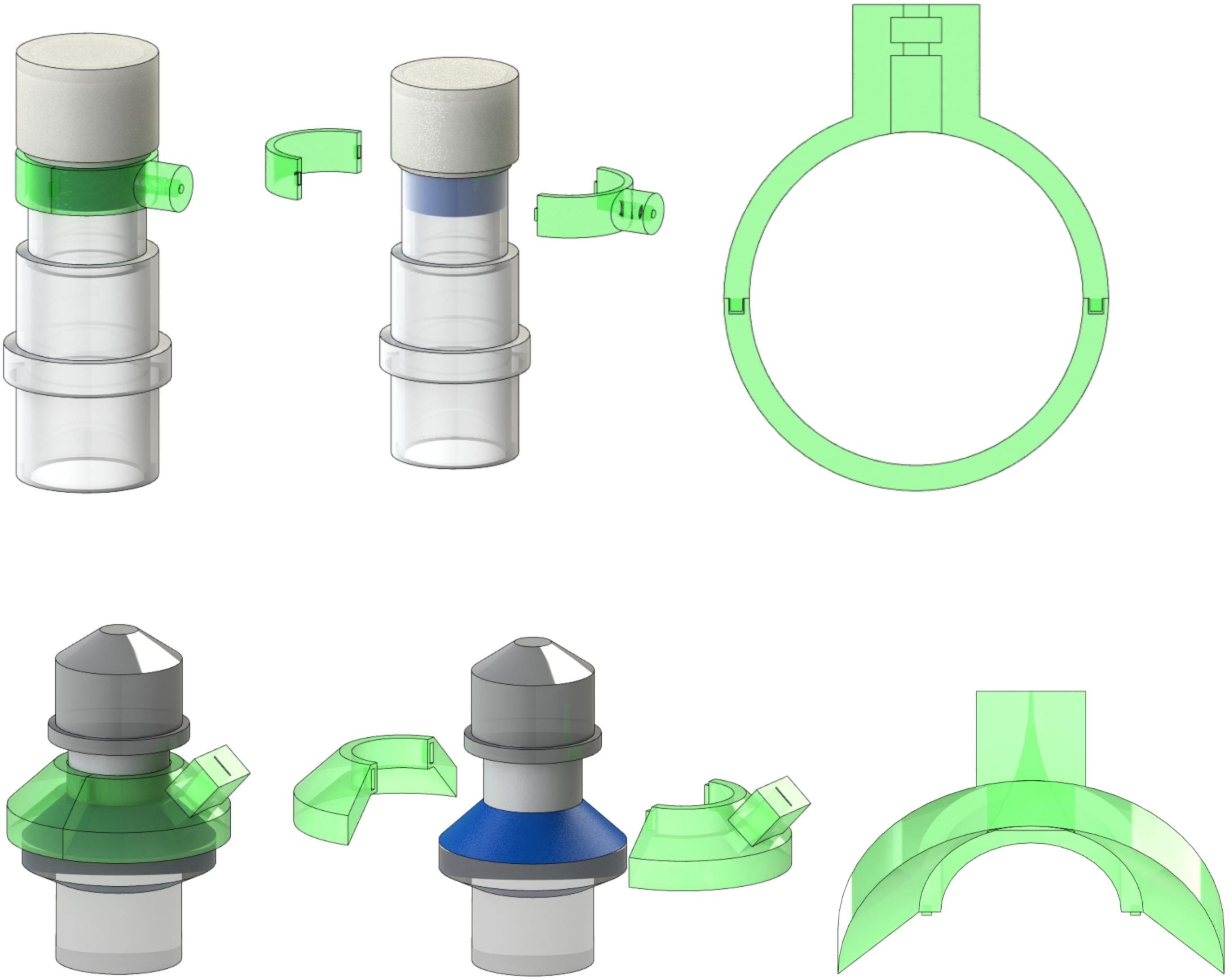

**Supplemental Figure 3.** Whistle ring designs for two types of common pressure release (PEEP) valves. These enable an auditory alarm when the PEEP valve actuates. This can be 3D printed and assembled onto standard PEEP valves with a 22mm inner diameter.

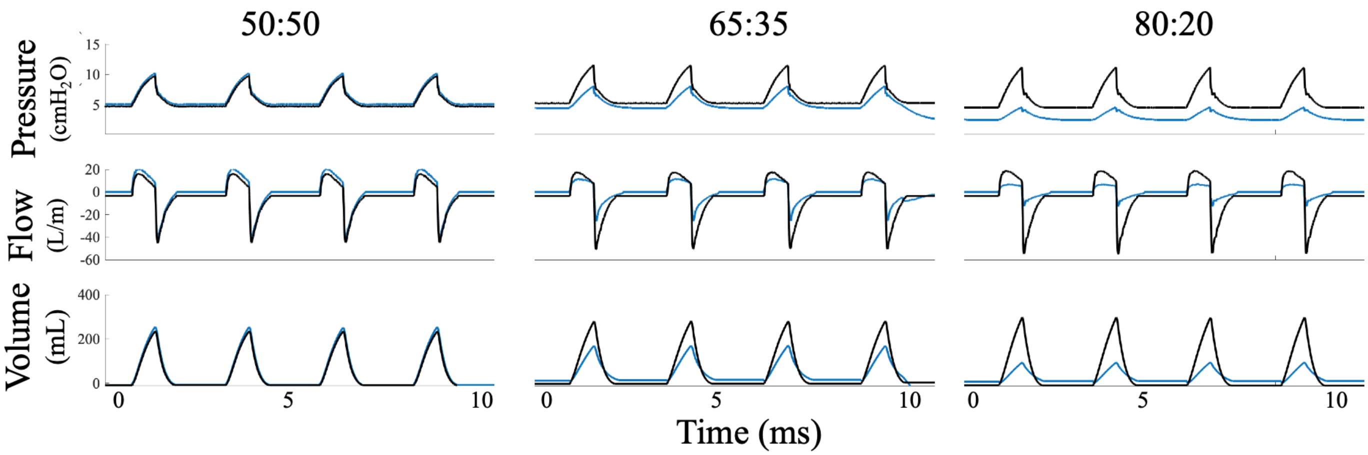


**Supplemental Figure 4. Variable Tidal Volume Production on the iSAVE employing an open circuit ventilator.** By adjusting the flow valve, various ratios of the ventilator's tidal volume of 400mL were delivered to the two artificial test lungs (capacity 1L). The range of ratios tested (50:50, 65:35, 80:20) demonstrate that the iSAVE can split the ventilator's tidal volume to patients of significantly different needs without creating unreasonable pressure of flow conditions.

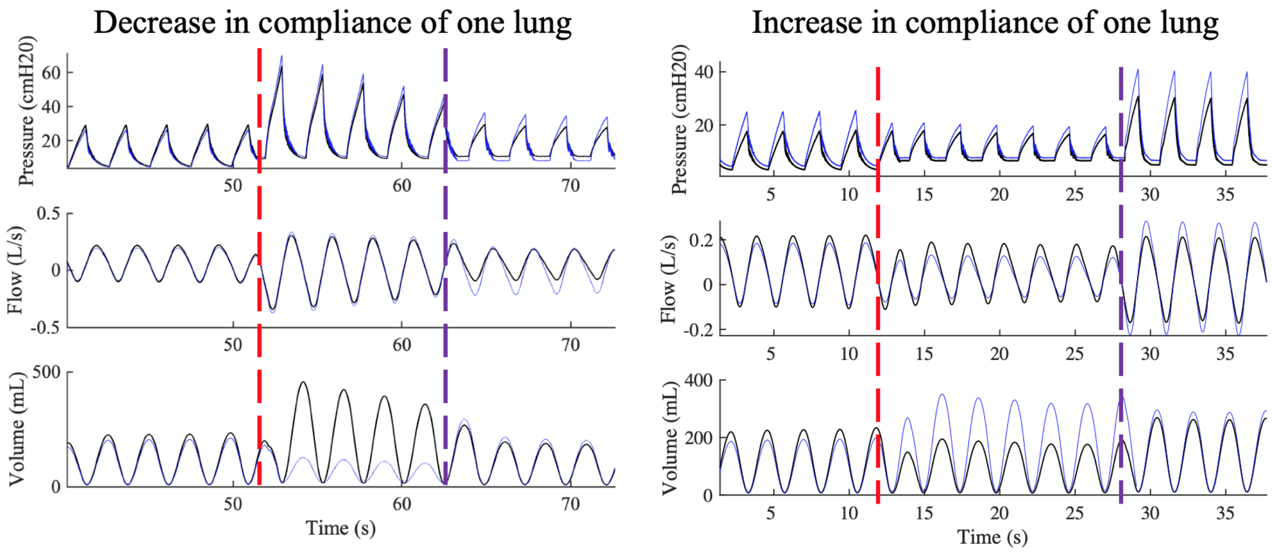


**Supplemental Figure 5. Accommodation of the iSAVE utilizing an open circuit ventilation to changes in compliance of one lung.** Initially, both lungs (C = 60 cm$H_2$0, R = Rp20) were supplied with $V_T$ = 200mL. (Left) The compliance of one lung was decreased (C = 30 cm$H_2$0) as indicated by the red dotted line, causing a shift in volume delivery to the other lung (black). Titration of the flow valve, indicated by the purple lines, restored flow to the baseline values. (Right) The compliance of one lung was increased (C = 120 cm$H_2$0, blue) as indicated by the red dotted line, causing a decrease in volume delivery to the other lung (black). Titration of the flow valve, indicated by the purple lines, restored flow to the baseline values.

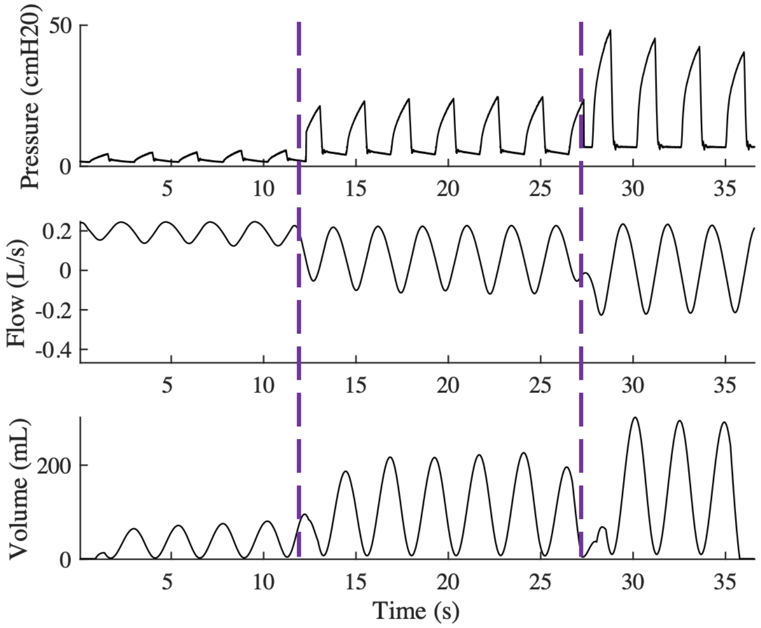


**Supplemental Figure 6. Titration of Valve at High Resistances.** The iSAVE was implemented using an open circuit ventilator to support two artificial lungs with high resistances (Rp50, C = 60 $cmH_20$). The valves were serially adjusted (as indicated by purple lines) to titrate flow. At the medium setting, 200mL was able to be delivered at a pressure of 22 $cmH_20$. The resistances simulated here are much higher resistance than what is expected in patients with COVID-19, even when comorbid with restrictive pathology. This test demonstrated the ability for the ventilator to supply multiple patients with sufficient volume, even when resistances are high.

**Supplemental Note 1: Ventilation Capacity**

Our testing thus far has provided a proof of concept for the iSAVE in a 1 ventilator : 2 patients set up ratio. Our testing demonstrated that ventilators' volume can be linearly divided to additional patients. Thus, the maximal capacity of this system is defined by:

$$Capacity = \frac{TV_{Ventilator}}{(TV_{Patient\ 1} + TV_{Patient\ 2} + TV_{Patient\ 3} + \cdots)}$$

Theoretically speaking, with most ICU ventilators able to provide up to 2500mL, it is estimated at least 6 patients could be simultaneously ventilated. However, a number of physical and practical challenges will limit the implementation. Dead space accumulated in the tubing cannot be greater than the lowest individual tidal volume required. Further, with the addition of more than two patients to one ventilator circuit, determining flow changes and titration will entail a more involved process, requiring monitors for each patient. In the case of *n* patients, data from *n-1* circuits would be required to make adjustments. If the supply of respiratory monitors is constrained, this certainly poses a limitation of the system, but can be overcome by sharing monitors and using them to perform periodic checks.
